# Human threat learning is associated with gut microbiota composition

**DOI:** 10.1101/2022.06.13.495985

**Authors:** Javiera P. Oyarzun, Thomas M. Kuntz, Yoann Stussi, Olivia T. Karaman, Sophia Vranos, Bridget L. Callaghan, Curtis Huttenhower, Joseph E. LeDoux, Elizabeth A. Phelps

## Abstract

Rodent studies have shown that the gut microbiota can influence threat and safety learning, which has been linked to anxiety phenotypes. In humans, it has been demonstrated that microbiota composition varies with anxiety disorders, but evidence showing an association with threat learning is lacking. Here, we tested whether individual variability in threat and safety learning was related to gut microbiota composition in healthy adults. We found that threat, but not safety learning varies with individuals’ microbiome composition. Our results provide evidence that the gut microbiota is associated with excitatory threat learning across species.

**Significance Statement:** Learning from threats and safety is a core mechanism of anxiety disorders, and studies in rodent models have shown that the gut microbiota can modulate such behaviors. Although previous literature on humans shows a relationship between emotional circuits and gut microbiota, the evidence linking learning and microbiota is lacking. In a Pavlovian threat conditioning paradigm, we show that patterns of gut microbiota composition in healthy humans relate to their patterns of threat learning, but not safety learning. Our findings suggest one mechanism by which the human gut microbiota is associated with anxiety-related behaviors.

## Introduction

Studies in rodents have demonstrated that the gut microbiota - the collective of all microorganisms that inhabit the host’s large intestine – can modulate hippocampal and amygdala-dependent learning as well as anxiety-like behaviors. For example, rodents treated with live bacteria show better memory recognition and decreased anxiety-like behaviors compared with non-treated or germ-free groups^1^. Furthermore, probiotic administration can have anxiolytic effects and reverse memory impairments observed after protocols of gut infection (through the administration of pathogenic bacteria) or chronic stress^2^.

One clinically relevant learning mechanism related to anxiety phenotypes is the ability to form associations with threat and safety outcomes. Studies in rodents have investigated threat and safety learning using Pavlovian threat acquisition and extinction tasks, respectively. Manipulations of gut microbiota through probiotics and heat-killed bacteria facilitate threat acquisition in rodents, as measured by enhanced defensive responses to threatful cues and contexts^3–5^. However, for extinction learning, results have been less consistent, with some showing extinction impairment or persistence of defensive^3,5^, others facilitation^6,7^, and still others no effect^4^ for safety learning after microbiota manipulations.

In humans, evidence linking threat and safety learning and microbiota composition is scarce and indirect. Both threat and safety learning^8^ and gut microbiota profiles^9^ are altered in anxiety patients compared to healthy individuals, and neuroimaging studies have shown an association between functional and structural brain circuits of threat processing and microbiota composition^10,11^. These previous studies suggest that there might be microbiota patterns associated with threat and safety learning in humans, but they do not directly assess this relationship. In the present study, we sought to test this possibility.

## Methods and Results

One hundred and twenty-seven healthy individuals (final sample size of 117 participants, see Extended Methods (EM) for inclusion criteria and Demographics, Table S1) underwent a Pavlovian threat conditioning paradigm while skin conductance responses (SCR) were recorded as a readout of defensive responses (Figure 1, see also EM).

**Figure 1.**
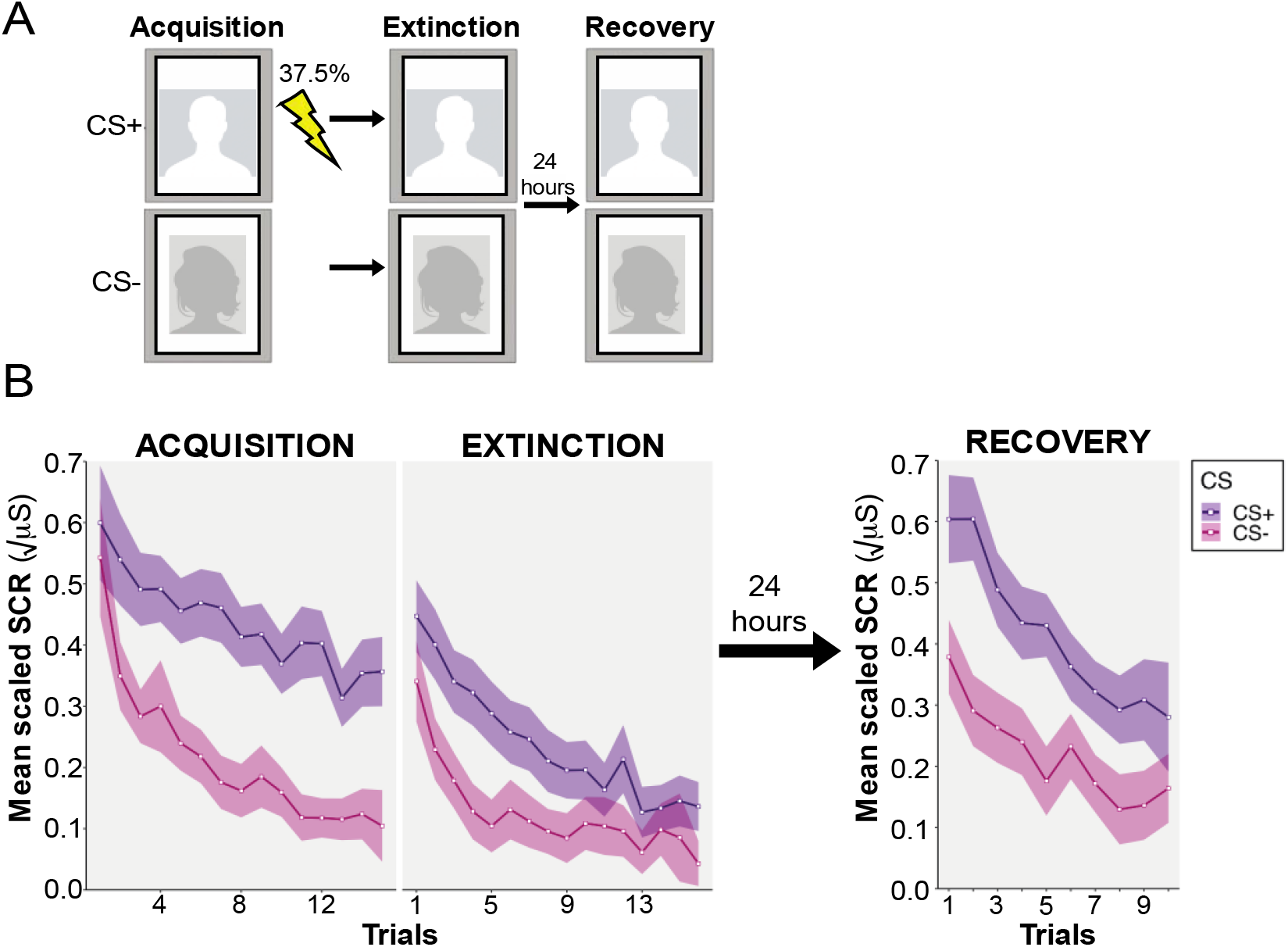
**A. Threat Conditioning Task.** *Acquisition:* participants were shown two fear faces. One picture, the conditioned stimulus (CS+ no-shock, 15 trials) was contingently paired with a mild electric shock to participants’ wrist (CS+ shock, 9 trials) according to a partial reinforcement schedule while the other was never paired with shock (CS-, 15 trials). *Extinction:* immediately following acquisition participants were presented with the CS+ and CS- (16 trials) without shock. *Recovery*: After 24 hours CS+ and CS-were presented without shock (10 trials each). **B**. Skin Conductance Responses during Threat Conditioning for non-shocked CS+ and CS-trials

**Figure 2.**
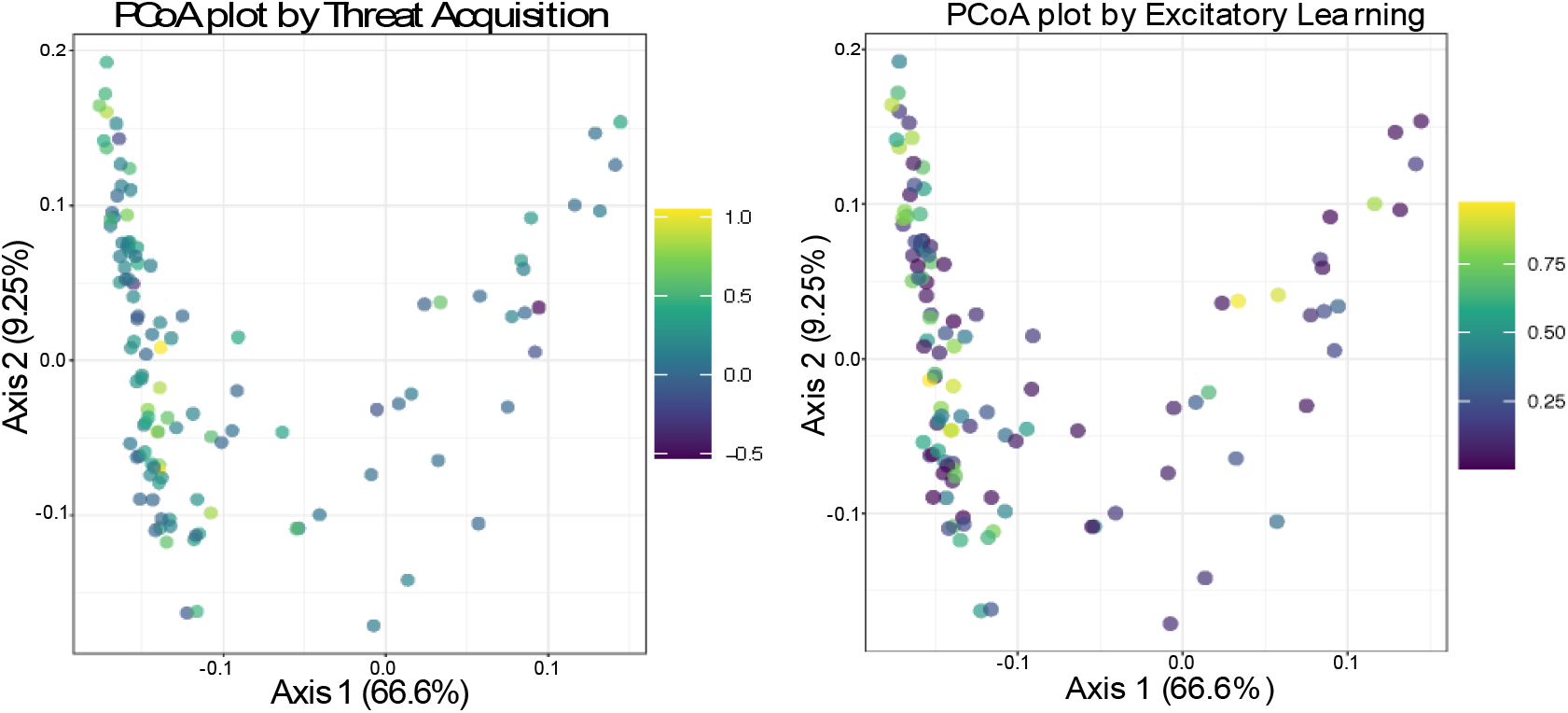
Principal Coordinate Analyses for Threat Acquisition and Excitatory Learning. Ordinations of all subjects’ microbial genera profiles by weighted UniFrac. While neither threat acquisition nor excitatory learning indices were associated with the two major axes of variation, as expected, both showed significant association with overall microbiome composition (Threat Acquisition: R^2^ = 0.045, *P* = 0.012; Excitatory Learning: R^2^ = 0.041, *P* = 0.009).

After the first session day or the next morning, participants provided a stool sample for 16S ribosomal RNA gene amplicon sequencing. Questionnaires were administered to assess psychological variables and factors that have been previously associated with gut microbiota composition, referred to as control variables: human milk feeding (*HMF*), mode of birth, exercise, current pet-cohabitation, childhood pet-cohabitation, and diet.

From the normalized SCR data, we calculated *threat acquisition, extinction learning*, and *recovery indexes* (see EM). Furthermore, since threat learning paradigms engage a combination of excitatory (i.e., updating due to presence of shock) and inhibitory (i.e., updating due to absence of shock) learning processes, we evaluated individuals’ bias to learn from the presence or absence of aversive outcomes. We fit a hybrid reinforcement model to the trial-by-trial SCR data from acquisition and extinction (see EM, Figure S1-S2) and extracted each participant’s excitatory and inhibitory learning rates.

We performed taxonomic profiling on our sequenced dataset and calculated associated ecological metrics from participants’ profiles. There was no association between gut microbiota alpha diversity (species Inverse Simpson index) and any threat learning parameters. Next, we performed a permutational multivariate analysis of variance (PERMANOVA, n=999 permutations) on genus level, weighted UniFrac beta diversities (see EM) in a multivariable model for each threat learning parameter with control variables. We found significant associations for the threat acquisition index (Weighted UniFrac R^2^ = 0.045, *P* = 0.012) and excitatory learning rates (Weighted UniFrac R^2^ = 0.041, *P* = 0.009) (see EM Tables S3a-b). These results suggest that participants that show a different index of threat acquisition and excitatory learning also show different patterns of microbiota composition.

To identify specific taxa associated with individuals’ threat acquisition index and excitatory learning, we used generalized linear models implemented in MaAsLin 2^12^ (see EM). Relative abundance of three taxa showed a significant relationship (FDR q<0.2) with participants’ acquisition index: UBA1819 (an unclassified *Faecalibacterium* taxon), *Tyzzerela*_3, and *Bacteroides*. However, these did not remain individually significant in a robustness analysis (see EM), and their linear associations were likely outlier-driven. To test for multivariable associations, we implemented a random forest classifier (see EM) across the same set of genus abundances. In agreement with our PERMANOVA results, the model significantly associated overall microbial community composition with threat acquisition index (R^2^ = 0.027, *P* = 0.022). The top 8 taxa included, *Butyricicoccus, Agathobacter, Alistipes, Phocea*, and V*eillonella*, in addition to the three taxa identified univariately above (Figure 3). This suggests that the combined relative abundance of various taxa is more important in threat acquisition than the abundance of any single taxon, and/or that inter-individual variation in specific microbial carriage makes univariate tests underpowered in this population.

**Figure 3.**
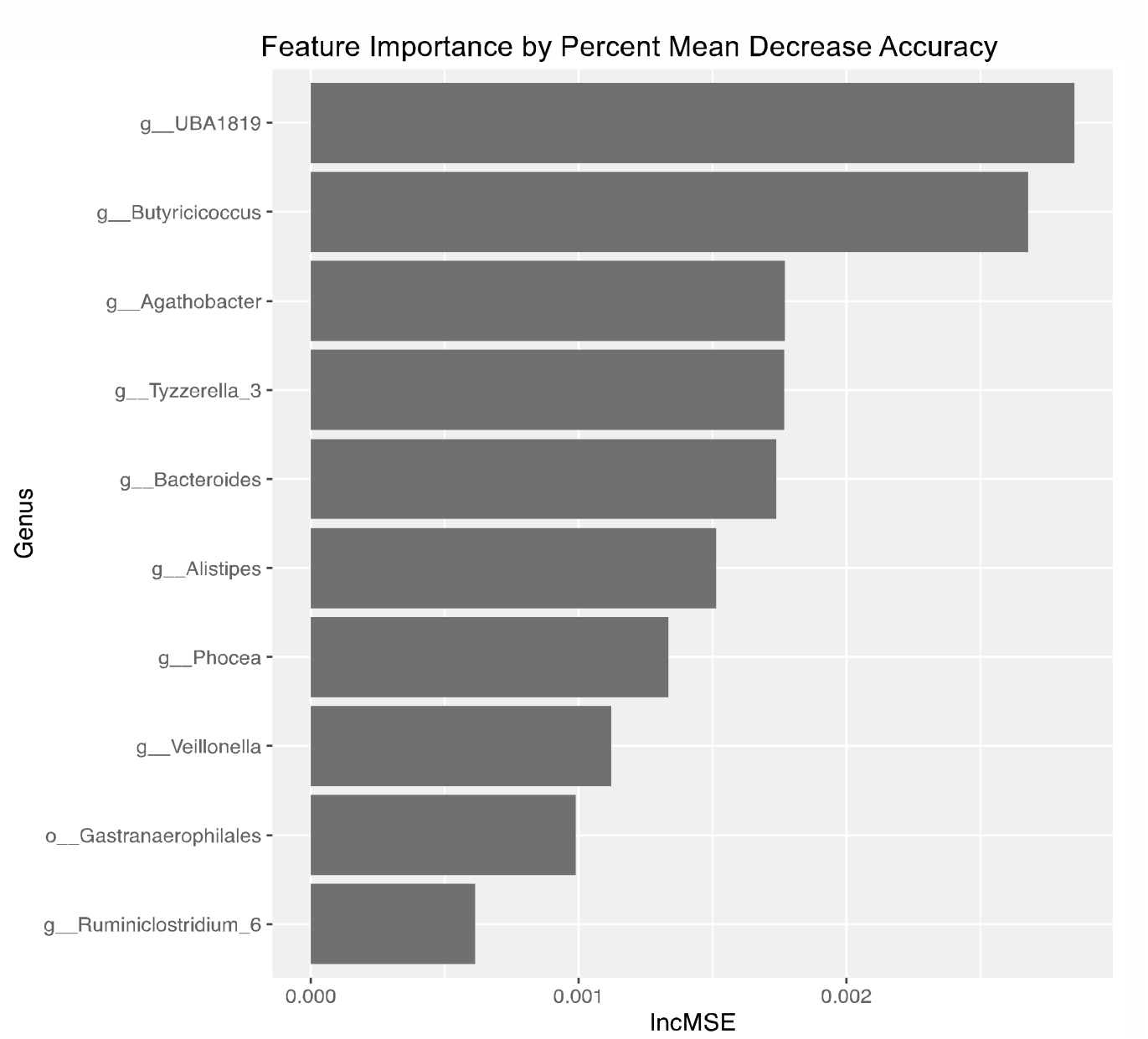
Feature Importance Table for Threat Acquisition. The table depicts the top 10 features predicting Threat Acquisition according to the Random Forest model (R^2^ = 0.027, *P* = 0.022). Importance scores were quantified by the percent increase in Mean Square Error when permuting features’ relative abundance values.

In contrast to rodent findings, no relationship between gut microbiota composition and extinction learning index, inhibitory learning, or recovery was observed (See EM; Figures S3b Extinction Learning, S3c Recovery Index, S3e Inhibitory Learning).

## Discussion

The association between the gut microbiota and learned defensive responses found here is in line with previous rodent research showing manipulations of the microbiota influence threat learning, as well as studies in humans showing that individuals with different microbiota compositions show differential amygdala structural and functional connectivity^10^, and other functional networks (dorsal anterior cingulate cortex, insula) ^11^ implicated in the processing and expression of threat learning^13^. Even though these previous studies were small in sample size, conducted in diverse populations (i.e., infants, adult women, obese adults, or patients with GI symptoms), and used different methods of characterizing gut microbiota composition^10^, these findings support the idea that the gut microbiota composition is associated with neural circuits implicated in threat acquisition and its expression.

Anxiety disorders have been associated with both enhanced acquisition of defensive responses to potential threats (excitatory learning) and impaired safety learning (inhibitory learning)^8^. Threat acquisition and extinction are mediated by overlapping but distinct, neural circuits^14^. The current results suggest that the relation between anxiety and microbiota composition in humans may depend more on enhanced learning of defensive responses than on the ventral prefrontal inhibitory circuits underlying extinction/inhibitory learning and recovery of defensive responses^13^.

Although theoretical pathways linking brain circuits of threat acquisition and gut microbiota have been suggested^2^ (e.g., enteric or vagal nervous signaling^3,15,16^, circulating microbial chemical products^7^, changes in nutrient processing and uptake, or joint differences due to developmental exposures), our results do not indicate specific mechanisms by which the gut microbiota might relate to threat circuits or even the specific direction of potential causality. In our dataset, no clear associations were found between individual taxon abundances and skin conductance responses; but rather broad microbiota patterns were indirectly associated. The associations found in our dataset may rely on microbial chemical products or functions shared by multiple taxa or external influences that affect both the microbiome and the threat acquisition. Alternatively, we may have lacked the power to find individual relationships with specific taxa. Larger study populations and the use of methods focused on microbiota functional profiling (metagenomics, transcriptomics, and metabolomics) could help identify potential mechanisms underlying current findings.

Even though our data cannot speak to an underlying mechanism, one possible biological pathway for this interaction is the influence of the gut microbiota on the host’s immune system and the effect of systemic inflammation on mood and anxiety symptoms^17^. For example, a recent meta-analysis showed that patients with psychiatric illnesses like depression and anxiety possessed an inflammatory phenotype associated with a low abundance of strict anaerobes in the gut, and a high abundance of potentially pro-inflammatory bacteria when compared with healthy individuals^18^. Although some of the taxa from the microbial community identified by our RF model have been previously associated with the absence of GI inflammation (i.e., *Butyricicoccus, Faecalibacterium, Bacteroides)*, there is a need for a larger cohort that expands the range of baseline microbial configurations – especially for low prevalence taxa – to confirm (or deny) the associations with defensive responses found here.

The present study found gut microbiota composition in humans is associated with threat acquisition and excitatory learning using two omnibus microbiome analysis techniques (PERMANOVA and Random Forest analyses). Although our sample only included healthy adults, the findings suggest that enhanced excitatory threat learning may be one factor contributing to the association between anxiety and gut microbiota. Emerging evidence linking the gut microbiome to mental health harbors the potential for innovative treatment approaches. However, it is necessary to determine the unique and modifiable factors that underlie this relationship. This finding represents a step in this direction by linking a specific learning component of an anxiety phenotype with overall microbiome composition in adult humans.

## Acknowledgments

We want to thank Dr. Anne-Catrin Uhlemann and Felix Rozenberg for their work during stool sample processing and sequencing, and Yasmine Elasmar, and Eugenia Zhukovsky for their help during data collection. We thank Dr. Jeremy Wilkinson for his support and insights during data analyses. This research has been supported by The Vulnerable Brain Project (vbp.life) to JEL and EAP, the James S. McDonnell Foundation to EAP, and the National Center for Advancing Translational Science (NCATS) from the National Institute of Health through the Grant Award Number TL1TR001447 to JPO.

## Extended Methods

### Participants

One hundred and twenty-seven participants (73 females, 20-50 years [M = 26.55, SD = 7.66]), Body Mass Index 16.42-37.97 [M = 24.48, SD = 4.37]) were recruited from the New York University (NYU) community via posted flyers and online cloud-based subject pool software (Sona Systems). The NYU Committee on Activities Involving Human Subjects approved all recruiting instruments and experimental protocols. Participants were paid $15 per hour and earned $50 for providing a stool sample.

#### Inclusion criteria

Eligible participants were between 20 and 50 years of age, proficient in English, had a normal or corrected vision, were not currently taking psychoactive medications, corticosteroids, antibiotics, or probiotics (see Table S1 for demographic details). Participants had no diagnosed medical disorders. Four participants reported being on an altered sleep cycle (working and studying during the night).

**Table S1.** Demographics and Control Variables

| N=127 | Mean | SD | Yes | No | NA |
| --- | --- | --- | --- | --- | --- |
| Age | 26.55 | 7.66 |  |  |  |
| BMI | 24.48 | 4.37 |  |  |  |
| Diet Mediterranean Adherence | 5.19 | 2.18 |  |  |  |
| Breastfeeding |  |  | 95 | 31 | 1 |
| C-section |  |  | 28 | 98 | 1 |
| Current pet co-habitation |  |  | 37 | 89 | 1 |
| Childhood pet |  |  | 96 | 30 | 1 |
| Exercise |  |  | 72 | 54 | 1 |
| Antibiotics under 3 years |  |  | 30 | 91 | 6 |

### Behavioral Task

#### Threat conditioning

Participants were tested on two consecutive days. Each participant was presented with two black-and-white images of fearful-expression faces from the Ekman pictures of facial affect^1^. Pictures were randomly selected out of a pool of 8 different male and female pictures. Pictures were presented over a gray background in a computer screen for 4s with a jittered inter-trial interval of 8–10s. Participants were instructed that one of the pictures – the CS+ – will be followed by an electric shock to the wrist, while the other picture – the CS-– will never be followed by a shock. They did not know beforehand, which picture was going to be paired with shocks. Electric shocks were delivered 37.5% of the times a CS+ was presented. To speed up threat conditioning, reinforcement was pseudo-randomized; the first stimuli were always reinforced, and there were never more than 3 consecutive reinforced CSs. Participants received a total of 9 reinforced CS+, 15 unreinforced CS+, and 15 CS-trials.

#### Extinction and recovery

During extinction and recovery sessions participants were exposed to the same pair of pictures, 16 and 10 times respectively. The same setup and temporal parameters of the acquisition phase were used in both sessions. No shocks were administered during these phases. The extinction session started 5 minutes after the end of the acquisition session and the recovery was conducted after 24 hours.

#### Shock stimulation and intensity calibration

The electric shocks (50 pulses/s) – here referred to as the unconditioned stimulus (US) – lasted 200ms and co-terminated with the respective picture presentation. Shocks were generated using an SD9 Square Pulse Stimulator (Grass Technologies) device and were delivered to participants’ dominant-hand wrist through two Ag/AgCl electrodes (EL503, Biopac) with conductance gel (GEL100, Biopac).

Each participant set the shock level at the beginning of the session. Participants received 5 shocks with increased intensity and were asked to rate their discomfort using a 0 to 9 scale (0 being no sensation to 9 being high intensity). To facilitate discomfort ratings, participants saw 10 cartoon faces with emotional expressions for each discomfort level. Calibration ended after the fifth shock when participants rated the shocks between intensity 6 and 8 and reported that the shock was “uncomfortable but not painful”. If participants rated the shock intensity outside the 6-8 range, up to 3 extra shocks would be administered to adjust the shock intensity to the 6-8 rate intensity window. The final shock level was fixed for the entire acquisition. Participants used a shock level between 10V and 65V (M = 38.41, SD = 12.23).

### Skin conductance responses, data acquisition, and scoring

Due to technical difficulties (i.e., electrodes disconnected from the participant, fire alarms went off, noise signal due to movements and talking during the session) and dropouts after the first day, skin conductance response (SCR) was acquired on 117 participants as an index of autonomic arousal. SCR signals were collected through two Ag/AgCl electrodes (EL507, Biopac) with NaCl gel (GEL101, Biopac) attached to the participants’ non-dominant palm. SCRs to stimuli were analyzed using Autonomate^2^ implemented in MATLAB 2017b (MathWorks). Autonomate scored SCR amplitudes (μSiemens) by looking at the highest peak generated between 0.5s and 5s after picture onset. If multiple peaks were present in this time window, the largest peak was scored. Reinforced trials were omitted. Autonomate-analyzed data were visually inspected to determine if the response detection was correct. When detection was incorrect, the correct peak of that trial was manually scored. Final trial scores were square-root-transformed and scaled according to each participant’s average SCR to the shocks acquired during the threat-acquisition phase. For this, peak SCRs to the 9 USs delivered, were detected during reinforced CS+ trials in a window between 4s and 8s after picture onset. SCRs were averaged and used to normalize the participant’s SCRs throughout the 3 sessions (acquisition, extinction, recovery).

#### Indexes calculation

Acquisition and extinction indexes were calculated by averaging the last 4 (unreinforced) CS+ and CS-trials. The recovery index was calculated by subtracting the extinction index to the average of the first 4 trials of the recovery session.

### Computational modeling of skin conductance response

We used reinforcement learning to characterize how variations in gut microbiota diversity relate to threat learning at a computational level (see refs^3,4^, for a similar modeling approach).

The learning parameters were estimated by fitting reinforcement learning models to the normalized (i.e., square-root-transformed and scaled) trial-by-trial SCR data during acquisition and extinction and Bayesian model selection was used to identify the model with the best fit to the data. The data from two participants were discarded from the computational analyses because their parameters could not be estimated due to a lack of SCR to all or some of the CS categories during the experiment, leaving a sample size of 115 participants (61 women; M_age_ = 26.34, SD_age_ = 7.56). The Hybrid model with dual learning rates was identified as the model most likely to have generated the observed data.

#### Hybrid model with dual learning rates

We adapted the Hybrid model proposed by Li et al.^5^ by implementing distinct learning rates for positive (i.e., when the shock is delivered and/or more than predicted; excitatory learning) and negative (i.e., when the shock is omitted or less than predicted; inhibitory learning) prediction errors^6^. This model combines the basic assumption of the Rescorla-Wagner model^7^ that learning is driven by prediction errors, while additionally incorporating the Pearce-Hall associability mechanism^8^. A cue’s associability corresponds to the extent to which the cue has been associated with surprise (i.e., prediction errors). It acts as a dynamic learning rate that controls the amount of future learning about the cue depending on its past reinforcement history. Specifically, associability accelerates learning to cues that are poor predictors of reinforcement, whereas it conversely decelerates learning to cues that reliably predict reinforcement. In concert with associability, excitatory and inhibitory learning rates govern the extent to which the organism learns about positive and negative prediction errors, respectively. The Hybrid model with dual learning rates posits that the expected value *V* of a given conditioned stimulus *j* at trial *t* + 1 is updated based on the sum of the current expected value *V*_*j*_ at trial *t* and the prediction error between *V*_*j*_ and the outcome *R* at trial *t*, weighted by (a) different learning rates for excitatory learning and inhibitory learning and (b) the associability *α* of the conditioned stimulus *j* at trial *t*, as follows:

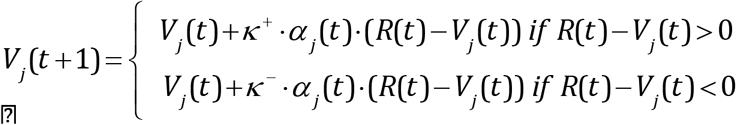

where the (static) excitatory learning rate *κ*^*+*^ and the (static) inhibitory learning rate *κ*^*–*^ are free parameters within the range [0, 1]. If the shock was delivered on the current trial *t, R*(*t*) = 1, else *R*(*t*) = 0. The associability *α*_*j*_ at trial *t* + 1 is, in turn, updated based on the sum of the absolute (i.e., unsigned) prediction error at trial *t* weighted by an associability weight, and the current associability *α*_*j*_ at trial *t* weighted by the complement to 1 of the associability weight:

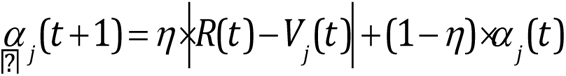

where the initial associability *α*_*0*_ and the associability weight *η* are free parameters within the range [0, 1]. The associability weight tracks participants’ relative sensitivity to past (*η* = 0) versus current (*η* = 1) prediction errors in the computation of associability.

#### Model and parameter fitting

The free parameters were fitted and optimized using maximum a posteriori estimation^6,9^. This estimation procedure consisted in finding the set of parameters that maximizes the likelihood of each participant’s trial-by-trial normalized SCRs to the CSs given the model, constrained by a regularizing prior. We constrained the free parameters using a Beta (1.2, 1.2) prior distribution slightly favoring values in the middle of the parameter range. We estimated the free parameters with the mfit toolbox (Gershman, 2016; https://github.com/sjgershm/mfit). To avoid local optima, we performed 20 random initializations to compute maximum likelihood estimates for each parameter. We modeled the loglikelihood of each trial’s normalized SCR LL_SCR(t)_ as a Gaussian distribution around a mean determined by the scaled expected value (or associability, or a combination of both expected value and associability) computed by the model, plus a constant β_**0**_and a variance parameter σ:

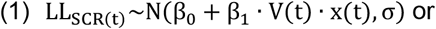

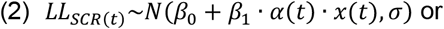

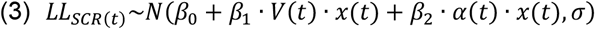

These are equivalent to linear regressions from the expected value time-series (1), the associability time-series (2), or the combination of both (3), to the normalized SCRs. We optimized the free parameters separately for each possible combination based on the trial-by-trial timeseries of values *V*(*t*), the trial-by-trial time-series of associabilities *α*(*t*), or the combination of both^5,10^. We excluded trials in which the US was delivered from the regressions onto the normalized SCRs to avoid possible contamination of the predictive response by responses related to the shocks^5,10^, but included them in the computation of the *V(t)* and *α*(*t*) timeseries.

As participants were expecting to receive electric shocks at the start of the threat conditioning session (because of the shock calibration procedure and the instructions), we set each CS initial expected value *V*_*0*_ to 0.5. We used a separate set of free parameters for each participant across acquisition and extinction trials.

#### Model comparison

We compared the Hybrid model with dual learning rates with alternative reinforcement learning models. Among the candidate models, we considered the classical Rescorla-Wagner model^11^, according to which the expected value of a given CS (*V*_*j*_(t+1)) is updated based on the sum of the current expected value (*V*_*j*_(t)) and the prediction error (*R*(*t*)- (*V*_*j*_(t)) weighted by a constant learning rate (*κ*), as follows:

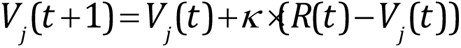

We also considered an adapted version of this model implementing separate learning rates for positive (*κ*^*+*^; excitatory learning rate) and negative (*κ*^*-*^; inhibitory learning rate) predictions errors:

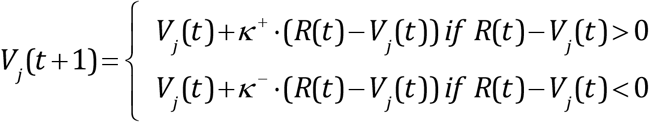

The last model we included was the Hybrid model^5^ incorporating a single (static) learning rate for both excitatory and inhibitory learning, in which the expected value (*V*_*j*_(*t*+1)) and associability (*α*_*j*_(*t*+1)) were updated as follows:

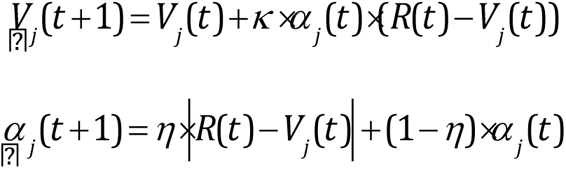

We performed model comparison using random-effects Bayesian model selection^12,13^. This procedure assumes that each participant is drawn from a single population distribution over models, which is estimated from the sample of model evidence values for each model^9,14^. The model evidence values were calculated using the Laplace approximation of the log marginal likelihood. We computed the protected exceedance probability (PXP) as a metric to compare the models. The PXP corresponds to the probability that a given model is more frequent in the population than all the other models under consideration while accounting for the possibility that some differences in model evidence may be due to chance. The Bayesian model selection procedure indicated that the Hybrid model with dual learning rates had a decisively higher PXP, suggesting that this model was more frequent in the population than the other models with a high probability (Figure S1).

**Figure S1.**
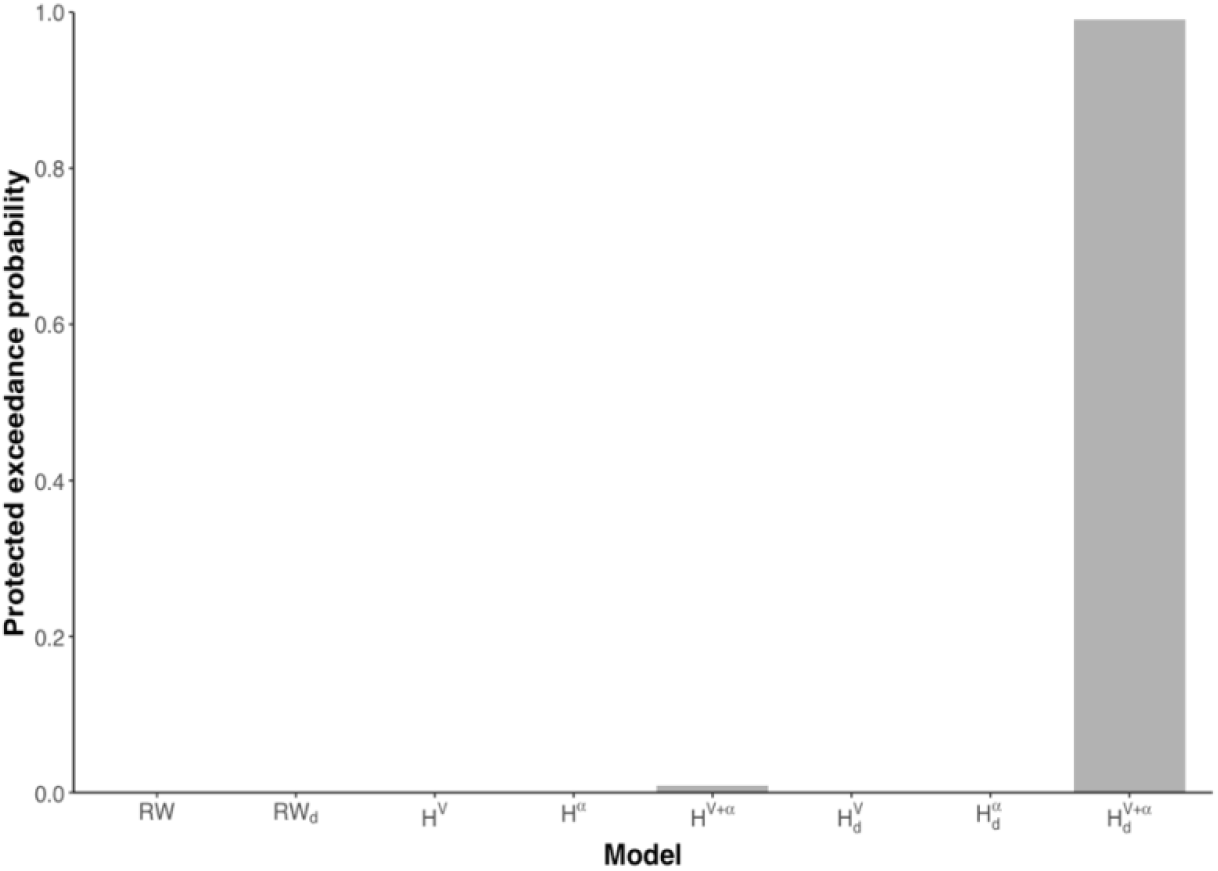
Bayesian model selection. The protected exceedance probability is reported for each model under consideration. RW = Rescorla-Wagner model, d = dual learning rates, H = hybrid model, V = expected values, *α =* associabilities.

#### Model and parameter recovery

We conducted a model recovery and a parameter recovery analysis to verify that the models can be correctly identified and the parameters recovered, respectively^***15***,***16***^. We ran 20 simulations that each generated predicted trial-by-trial expected values (and associabilities for the Hybrid models) for 115 synthetic participants with each of the models under consideration (see ref^***17***^, for a similar approach). For each simulation, we randomly sampled all parameters from a Beta (1.2, 1.2) distribution. The structure and properties of the task were identical to the 115 instances the actual participants experienced during the study. The models were subsequently fitted to the simulated data to (a) examine whether the model that generated the simulated data was identified as the most likely model across simulations (i.e., associated with the highest averaged PXP) and at the individual simulation level (i.e., associated with the highest PXP for each simulation) using Bayesian model selection, and (b) estimate the free parameters to assess whether the recovered parameter values were tightly correlated with the simulated parameters. For the Hybrid model and the Hybrid model with dual learning rates, the free parameters were estimated based on the combination of both values and associability timeseries. The model recovery analysis reflected that all the models were correctly identified in all the simulations. Linear regressions between the simulated and estimated parameters showed that regression intercepts (***β***_***0***_ values) were equivalent or close to 0 (all ***β***_***0***_s < 0.001), and the regression slopes (***β***_***1***_ values) equivalent or close to 1 (all ***β***_***1***_s > 0.998) and statistically significant (all ***P***s < 0.001). This indicates that all the Data from 2300 synthetic participants (20 simulations × 115 individuals) were simulated with the model. The four free parameters per participant were then regressed against the true parameters used for simulating the data. Results show excellent identifiability with regression intercepts (*β*_*0*_) close to 0, and regression slopes (*β*_*1*_) close to 1 and statistically significant (all *P*s < 0.001). Each dot represents a synthetic participant. The red lines represent the best linear fits. The gray densities represent the probability distribution used to sample the parameters. (B) Confusion matrices representing the Pearson correlations (top) and explained variances (bottom) between parameters at the individual simulation level. The parameters were estimated over 115-participants simulations and averaged over the 20 simulations. Diagonal: correlations between simulated and estimated parameters. Off-diagonal: cross-correlations between estimated parameters. *κ*^+^ = excitatory learning rate, *κ*^−^ = inhibitory learning rate, *α*_*0*_ = initial associability, *η* = associability weight. parameters were successfully recoverable across all the models. The estimated parameters from the Hybrid model with dual learning rates showed excellent recovery (see Figure S2A). At the individual simulation level, the Pearson correlations between the simulated and estimated parameters over 115 synthetic participants were high for all the models (RW: all averaged ***r***s = 1, all averaged R^2^ = 1; RW_d_: all averaged ***r***s = 1, all averaged R^2^ = 1; H: all averaged ***r***s =1, all averaged R^2^ = 1; H_d_: all averaged ***r***s > 0.99, all averaged R^2^ > 0.99). No cross-correlation was observed between estimated parameters (RW_d_: all averaged R^2^ < 0.02; H: all averaged R^2^ < 0.02; H_d_: averaged R^2^ < 0.01). The averaged Pearson correlations and explained variance between the simulated and estimated parameters at the single simulation level for the hybrid model with dual learning rates are shown in Figure S2B.

**Figure S2.**
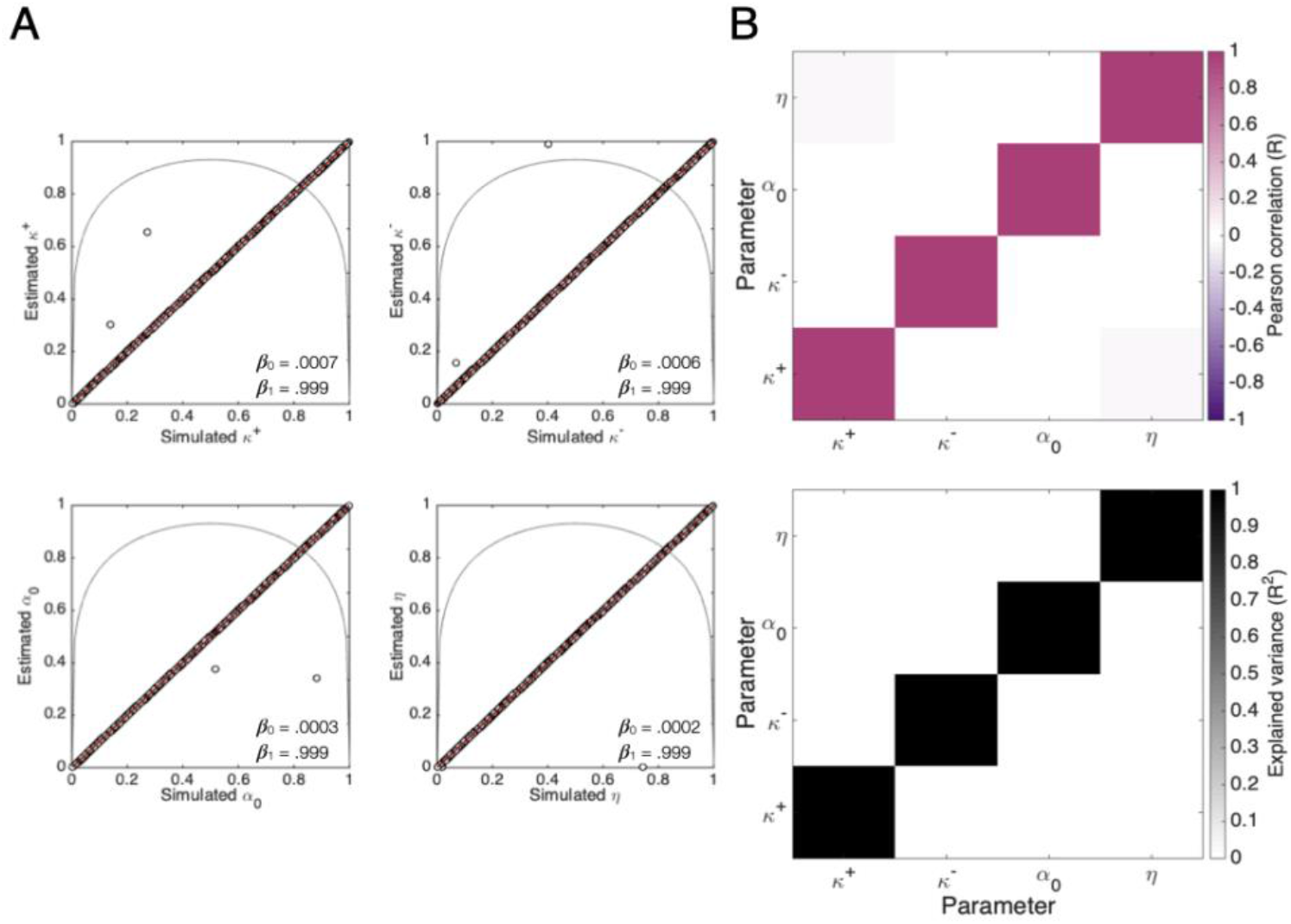
Parameter recovery analysis for the hybrid model with dual learning rates. (A) Data from 2300 synthetic participants (20 simulations × 115 individuals) were simulated with the model. The four free parameters per participant were then regressed against the true parameters used for simulating the data. Results show excellent identifiability with regression intercepts (β_0_) close to 0, and regression slopes (β_1_) close to 1 and statistically significant (all Ps < 0.001). Each dot represents a synthetic participant. The red lines represent the best linear fits. The gray densities represent the probability distribution used to sample the parameters. (B) Confusion matrices representing the Pearson correlations (top) and explained variances (bottom) between parameters at the individual simulation level. The parameters were estimated over 115-participants simulations and averaged over the 20 simulations. Diagonal: correlations between simulated and estimated parameters. Off-diagonal: cross-correlations between estimated parameters. ?^+^ = excitatory learning rate, κ^−^ = inhibitory learning rate, α _0_ = initial associability, η = associability weight.

#### Relationship between modeled learning signals and participants’ normalized skin conductance responses

We further examined the extent to which modeled expected value and associability signals from the Hybrid model with dual learning rates predicted participants’ trial-by-trial normalized SCRs^***5***,***18***^. To do so, we performed a multiple linear regression in which the expected value and associability timeseries generated with the individual parameter estimates from the model and averaged across all participants was regressed onto the averaged trial-by-trial normalized SCRs, excluding trials in which the US was delivered. This analysis indicated that expected value and associability signals explained a statistically significant amount of variance of the trial-by-trial normalized SCRs (R^2^ = .865, 90% CI [.794, .907], adjusted R^2^ = .861, *F*(1, 59) = 189.70, *P* < 0.001). Expected value signals predicted the trial-by-trial normalized SCRs, *b* = 0.720, 95% CI [0.304, 1.135], *β* = 0.646, *t*(59) = 3.47, *P* < 0.001, which was not the case for associability signals, *b* = 0.309, 95% CI [-0.087, 0.705], *β* = 0.291, *t*(59) = 1.56, *P* = 0.124.

#### Skin conductance responses between sex

Since previous literature has found interaction effects between behavior and microbiota composition in model organisms, we first assessed behavior across sex. Behavioral data were analyzed using R^19^. Shapiro-Wilk Normality test was used to assess the normality of the variables. Behavioral comparisons between sex groups were analyzed using the Wilcox test for non-normal data and an unpaired *t*-test for normally distributed data. No differences were found in behavior between sex groups. Follow-up analyzes were then continued using the gender-mix sample.

**Table S2.** Behavioral comparisons between sex.

|  | *P*(Shapiro-Wilk) | W (Wilcox) | *P*(Wilcox) | t(t-test) | *P*(t) |
| --- | --- | --- | --- | --- | --- |
| Acquisition Index | 0.000 | 1866 | 0.36 |  |  |
| Extinction Index | 1.19e-09 | 1423 | 0.16 |  |  |
| Recovery Index | 0.37 |  |  | 0.48 | 0.63 |
| Excitatory Learning ($\kappa+$) | 1.60e-07 | 1604 | 0.81 | | |
| Inhibitory Learning ($\kappa-$) | 4.25e-10 | 1774 | 0.47 | | |
| Initial Associability ($\alpha_0$) | 1.93e-05 | 1506 | 0.43 | | |
| Associability Weight ($\eta$) | 2.71e-08 | 1708 | 0.73 | | |

### Gut microbiota collection, processing, and analyses

#### Collection and DNA processing

At the end of the first experimental day, participants were provided with an OMR-200 OMNIgene-Gut stool collecting kit (DNA Genotek) and an OM-AC1 toilet accessory. Participants watched a video with the instructions about how to use the collecting kit at home and received further verbal instructions from a research assistant who answered any questions. Participants were instructed to collect a stool sample during the next bowel movement and bring it to the lab on the following day. Stool samples were stored at −20°C after receipt. Samples were then processed and sequenced in one batch at the Columbia University Microbiome Core (CUMC). DNA was extracted using the MagAttract PowerSoil kit (Mo-Bio), and 16S rRNA amplicon sequencing of the V3-V4 region with the 341F/805R primer pair^20^ was performed on the Illumina MiSeq 2×300 platform following the manufacturer’s recommended best practices. Resulting FASTQ files were bioinformatically processed through the bioBakery AnADAMA2 0.90 workflows^21^. Sequences were demultiplexed with ea-utils 1.1.2^22^. Cutadapt 3.0^23^ was used to remove remaining primer and adapter sequences. DADA2 1.14.1^24^ was used to denoise, filter, trim, merge, remove chimeras, and taxonomically classify the data, with default parameters other than a truncation length of 265bp for forward reads and 225bp for reverse as reads after these points have substantially lower average quality scores and interfered with read pair merging. Phylogenetic trees were constructed after the alignment of sequences using Clustal Omega 1.2.4^25^ with FastTree2.1.10^26^. ASVs were taxonomically assigned using the SILVA 132^27^ reference database. Features were filtered requiring at least 0.01% relative abundance in 10% of samples for all downstream analyses.

#### Beta diversity

Beta diversity was quantified by weighted UniFrac distance {16332807}. The variance explained (R^2^) and statistical significance for each behavior variable in multivariable models with control variables was calculated using Permutational Multivariable Analysis of Variance (PERMANOVA) tests (Tables S3a-g). To visualize the community compositions, weighted UniFrac distances were ordinated by Principal Coordinates Analysis (PCoA). Calculations were performed with the vegan 2.5-7^28^ package and visualizations were created using ggplot2 3.3.5, both in the R 4.1.0 computing enviroment^29^.

#### Feature associations

To identify specific taxa associated with individuals’ threat acquisition index and excitatory learning rates, we used general linear models as implemented in MaAsLin2 1.73^30^. MaAsLin2 1.73 calculates each feature’s associations with metadata with multiple comparison adjustments across all models. Filtered, normalized, relative abundance data is log-transformed before testing with a pseudo count of half of the lowest non-zero relative.

Results are adjusted using the Benjamini and Hochberg method and FDR *P*-values of 0.20 or lower are reported as significant. Univariable analyses showed a relationship between acquisition index and 3 genera: *Bacteroides* (coef=.080, sdterr=.0266, pval= .003, qval=.174), *Tyzzerella_3* (coef=.101, sdterr=.0266, pval= .002, qval=.174), and *UBA1819* (coef=.142, sdterr=.0525, pval= .007, qval=.201). These associations however did not remain individually significant in a robustness analysis, removing points 1.5 times the interquartile range (IQR) from the top quartile of the transformed relative abundances (after removing zeros) and 1.5 times the IQR from both sides of the acquisition index, suggesting that their linear associations were likely outlier driven.

#### Random Forest classifier

To investigate the predictive performance of the microbiome on outcome variables, random forest (RF) models^31^ were built using the R package randomForest 4.7^32^ including assessment of importance of individual predictors. The model was built on the transformed relative abundances as above, using 10,001 trees with otherwise default parameters. Model significance was determined via permutation (n = 1000) using the rfUtilities 2.1 package^33^.

Only the model for threat acquisition was significant (R^2^ = 0.027, *P* = 0.022). Importance scores were calculated by percent increase in Mean Squared Error (Mean Decrease Accuracy) and total decrease in node impurities, measured by the Gini Index from splitting on the variable, averaged over all trees (Mean Decrease Gini) and the values for the top 10 features were plotted (Figure 3, main manuscript)

**Table S3a.** Weighted UniFrac PERMANOVA on **Threat Acquisition Index**

|  | Df | SumOfSqs | R2 | F | Pr(>F) |
| --- | --- | --- | --- | --- | --- |
| **Threat Acquisition Index** | **1** | **0.327** | **0.045** | **5.634** | **0.012** |
| Age | 1 | 0.044 | 0.006 | 0.757 | 0.420 |
| BMI | 1 | 0.049 | 0.007 | 0.851 | 0.420 |
| **Diet** | **1** | **0.185** | **0.026** | **3.185** | **0.045** |
| Sex | 1 | 0.029 | 0.004 | 0.499 | 0.603 |
| Exercise | 1 | 0.024 | 0.003 | 0.411 | 0.690 |
| Human Milk Feeding | 1 | 0.144 | 0.020 | 2.485 | 0.101 |
| C-Section | 1 | 0.045 | 0.006 | 0.769 | 0.450 |
| Pet-current | 1 | 0.031 | 0.004 | 0.536 | 0.569 |
| Pet-childhood | 1 | 0.135 | 0.019 | 2.317 | 0.101 |
| Residual | 105 | 6.099 | 0.843 | NA | NA |
| Total | 115 | 7.239 | 1.00 | NA | NA |

**Table S3b.**
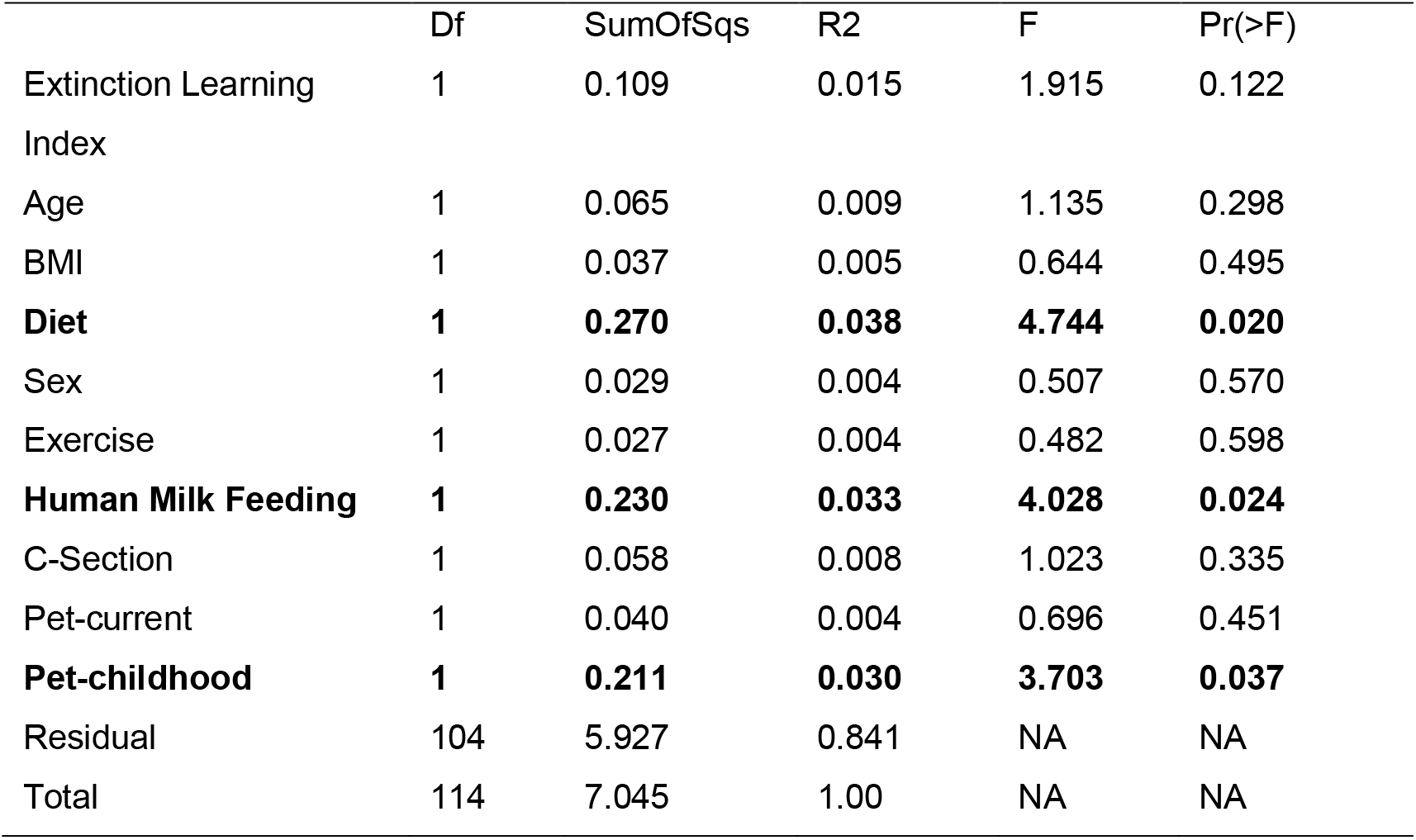
Weighted UniFrac PERMANOVA on Extinction Learning Index

|  | Df | SumOfSqs | R2 | F | Pr(>F) |
| --- | --- | --- | --- | --- | --- |
| **Extinction Learning Index** | **1** | **0.109** | **0.015** | **1.915** | **0.122** |
| Age | 1 | 0.065 | 0.009 | 1.135 | 0.298 |
| BMI | 1 | 0.037 | 0.005 | 0.644 | 0.495 |
| **Diet** | **1** | **0.270** | **0.038** | **4.744** | **0.020** |
| Sex | 1 | 0.029 | 0.004 | 0.507 | 0.570 |
| Exercise | 1 | 0.027 | 0.004 | 0.482 | 0.598 |
| **Human Milk Feeding** | **1** | **0.230** | **0.033** | **4.028** | **0.024** |
| C-Section | 1 | 0.058 | 0.008 | 1.023 | 0.335 |
| Pet-current | 1 | 0.040 | 0.004 | 0.696 | 0.451 |
| **Pet-childhood** | **1** | **0.211** | **0.030** | **3.703** | **0.037** |
| Residual | 104 | 5.927 | 0.841 | NA | NA |
| Total | 114 | 7.045 | 1.00 | NA | NA |

**Table S3c.** Weighted UniFrac PERMANOVA on **Recovery Index**

|  | Df | SumOfSqs | R2 | F | Pr(>F) |
| --- | --- | --- | --- | --- | --- |
| Recovery Index | 1 | 0.031 | 0.005 | 0.577 | 0.535 |
| Age | 1 | 0.077 | 0.012 | 1.431 | 0.192 |
| BMI | 1 | 0.023 | 0.004 | 0.43 | 0.628 |
| **Diet** | **1** | **0.278** | **0.043** | **5.166** | **0.010** |
| Sex | 1 | 0.028 | 0.004 | 0.521 | 0.584 |
| Exercise | 1 | 0.025 | 0.004 | 0.471 | 0.598 |
| **Human Milk Feeding** | **1** | **0.225** | **0.035** | **4.194** | **0.013** |
| C-Section | 1 | 0.034 | 0.005 | 0.639 | 0.506 |
| Pet-current | 1 | 0.050 | 0.008 | 0.937 | 0.337 |
| **Pet-childhood** | **1** | **0.264** | **0.041** | **4.923** | **0.015** |
| Residual | 101 | 5.426 | 0.839 | NA | NA |
| Total | 111 | 6.466 | 1.00 | NA | NA |

**Table S3d.** Weighted UniFrac PERMANOVA on **Excitatory Learning Rates**

|  | Df | SumOfSqs | R2 | F | Pr(>F) |
| --- | --- | --- | --- | --- | --- |
| **Excitatory Learning** | **1** | **0.297** | **0.041** | **5.075** | **0.009** |
| Age | 1 | 0.119 | 0.017 | 2.023 | 0.121 |
| BMI | 1 | 0.037 | 0.005 | 0.638 | 0.468 |
| Diet | 1 | 0.217 | 0.030 | 3.698 | 0.039 |
| Sex | 1 | 0.025 | 0.003 | 0.425 | 0.659 |
| Exercise | 1 | 0.020 | 0.003 | 0.336 | 0.716 |
| **Human Milk Feeding** | **1** | **0.201** | **0.028** | **3.437** | **0.033** |
| C-Section | 1 | 0.042 | 0.006 | 0.719 | 0.460 |
| Pet-current | 1 | 0.022 | 0.003 | 0.377 | 0.661 |
| **Pet-childhood** | **1** | **0.223** | **0.031** | **3.797** | **0.039** |
| Residual | 103 | 6.036 | 0.842 | NA | NA |
| Total | 113 | 7.167 | 1.000 | NA | NA |

**Table S3e.** Weighted UniFrac PERMANOVA on **Inhibitory Learning Rates**

|  | Df | SumOfSqs | R2 | F | Pr(>F) |
| --- | --- | --- | --- | --- | --- |
| Inhibitory Learning | 1 | 0.163 | 0.023 | 2.722 | 0.096 |
| Age | 1 | 0.100 | 0.014 | 1.669 | 0.156 |
| BMI | 1 | 0.023 | 0.003 | 0.391 | 0.681 |
| Diet | 1 | 0.216 | 0.030 | 3.607 | 0.042 |
| Sex | 1 | 0.027 | 0.004 | 0.456 | 0.611 |
| Exercise | 1 | 0.021 | 0.003 | 0.352 | 0.696 |
| Human Milk Feeding | 1 | 0.202 | 0.028 | 3.379 | 0.033 |
| C-Section | 1 | 0.055 | 0.008 | 0.915 | 0.385 |
| Pet-current | 1 | 0.025 | 0.003 | 0.417 | 0.630 |
| **Pet-childhood** | **1** | **0.267** | **0.037** | **4.462** | **0.025** |
| Residual | 103 | 6.171 | 0.861 | NA | NA |
| Total | 113 | 7.167 | 1.00 | NA | NA |

**Table S3f.** Weighted UniFrac PERMANOVA on **Initial Associability**

|  | Df | SumOfSqs | R2 | F | Pr(>F) |
| --- | --- | --- | --- | --- | --- |
| Initial Associability | 1 | 0.026 | 0.004 | 0.422 | 0.644 |
| Age | 1 | 0.095 | 0.013 | 1.544 | 0.181 |
| BMI | 1 | 0.029 | 0.004 | 0.478 | 0.602 |
| Diet | 1 | 0.204 | 0.028 | 3.332 | 0.046 |
| Sex | 1 | 0.021 | 0.003 | 0.344 | 0.723 |
| Exercise | 1 | 0.021 | 0.003 | 0.350 | 0.695 |
| Human Milk Feeding | 1 | 0.222 | 0.031 | 3.622 | 0.030 |
| C-Section | 1 | 0.067 | 0.009 | 1.100 | 0.319 |
| Pet-current | 1 | 0.021 | 0.003 | 0.344 | 0.699 |
| **Pet-childhood** | **1** | **0.212** | **0.030** | **3.459** | **0.049** |
| Residual | 103 | 6.308 | 0.880 | NA | NA |
| Total | 113 | 7.167 | 1.00 | NA | NA |

**Table S3g.** Weighted UniFrac PERMANOVA on **Associability Weight**

|  | Df | SumOfSqs | R2 | F | Pr(>F) |
| --- | --- | --- | --- | --- | --- |
| Associability Weight | 1 | 0.063 | 0.009 | 1.027 | 0.324 |
| Age | 1 | 0.095 | 0.013 | 1.560 | 0.174 |
| BMI | 1 | 0.020 | 0.003 | 0.326 | 0.759 |
| Diet | 1 | 0.192 | 0.027 | 3.149 | 0.058 |
| Sex | 1 | 0.025 | 0.003 | 0.408 | 0.665 |
| Exercise | 1 | 0.021 | 0.003 | 0.348 | 0.692 |
| Human Milk Feeding | 1 | 0.172 | 0.024 | 2.819 | 0.061 |
| C-Section | 1 | 0.063 | 0.009 | 1.033 | 0.339 |
| Pet-current | 1 | 0.022 | 0.003 | 0.367 | 0.667 |
| **Pet-childhood** | **1** | **0.224** | **0.031** | **3.672** | **0.042** |
| Residual | 103 | 6.271 | 0.875 | NA | NA |
| Total | 113 | 7.167 | 1.000 | NA | NA |

## Notes

### Competing Interest Statement

The authors have declared no competing interest.

